# The temperature sensitivity of arboviral disease extrinsic incubation periods: a systematic review

**DOI:** 10.1101/2025.08.08.669272

**Authors:** Anna-Maria Hartner, Tim Herath, Luis Roger Esquivel Gomez, Nils Körber, Juliane Pfeil, Jean-Baptiste Escudié, Lothar Wieler, Christopher Irrgang

## Abstract

**Background:** Arboviral diseases are an increasing global public health concern, driven by both human and environmental factors. A key parameter shaping transmission is the extrinsic incubation period (EIP)—the time between a mosquito acquiring a virus and becoming infectious—which is strongly influenced by temperature. However, the temperature–EIP relationship remains poorly characterized across arboviruses and mosquito species.

**Methods:** We conducted the first systematic review of laboratory studies evaluating temperature effects on EIP-related outcomes across all reported mosquito–arbovirus pairings. We searched three databases and extracted data on transmission efficiency (TE), applying linear regression models adjusted for diurnal temperature range and viral dose. Studies where TE could not be extracted were synthesized narratively.

**Results:** Our synthesis included 60 studies covering 17 arboviral diseases and 20 mosquito species. We found substantial heterogeneity in temperature effects on TE. While CHIKV, ZIKV, DENV, and WNV generally showed increased TE and shorter EIP at higher temperatures, many viruses—such as SINV, USUV, and BATV—exhibited no clear trends, often due to limited data and small sample sizes. Even across different vectors of the same virus, findings varied widely, reflecting both biological differences and inconsistent experimental designs.

**Conclusions:** These findings reveal major gaps in our understanding of climate-sensitive arbovirus transmission. Standardization of vector competence experiments and expanded research on neglected viruses are urgently needed. Environmental cofactors beyond temperature—such as humidity and variability—should also be incorporated to improve modelling and support climate-resilient intervention strategies.

**Author summary:** The burden of disease caused by arboviruses is increasing globally, largely driven by rapid urbanization, climate change, and global mobility. The transmission dynamics of arboviruses are modulated by a parameter known as the extrinsic incubation period, which is driven by environmental factors, especially temperature. However, the extent and direction of this effect has not been well characterized for all viruses, as data is limited and has not been synthesized for many neglected tropical diseases. Our study is the first systematic review to examine the existing evidence of laboratory studies across all studied mosquito species and virus pairings and to provide a comprehensive dataset for future modelling studies. We find that there is significant heterogeneity, not only between viruses, but also within individual vector-virus combinations, and that many arboviruses show no temperature effect. Based on these results, we propose that existing research efforts be standardized, and further research is conducted on emerging and re-emerging arboviruses for better intervention strategies and modelling predictions.

## Introduction

Arboviruses, transmitted by haematophagous arthropods such as mosquitoes, constitute a formidable challenge to global public health due to their widespread prevalence and capacity to cause severe disease. It is estimated that pathogens such as Dengue, Zika, Chikungunya, and West Nile virus collectively infect millions of people annually, with dengue alone accounting for over three million cases and 1,400 deaths since the beginning of 2025 [1]. These diseases impose substantial morbidity, mortality, and economic costs, particularly in tropical and subtropical regions [2, 3]. Though there have been increasing efforts in vector control and disease prevention, such as through vaccine development or the introduction of Wolbachia, these still pose significant challenges [4, 5]. For instance, the Dengue vaccine is currently recommended only for individuals with prior confirmed Dengue infection, which restricts its use in many endemic regions [4]. Additional efforts to expand the drugs, diagnostics, and vaccines available for arbovirus infections, are limited, and these diseases still remain difficult to control, with treatment access limited in rural areas, and existing treatments themselves often limited to symptom control [4]. As a result of these limitations, arboviral infections remain widespread globally. Compounding the issue, the frequency and scale of arbovirus outbreaks have been increasing due to a confluence of factors such as rapid urbanization, growing global mobility, and environmental changes [6, 7].

Climate change especially has been a major implicated factor in the expansion of the incidence and distribution of mosquito-borne arboviruses [8]. In this context, environmental factors have become increasingly recognized for their pivotal role in shaping arbovirus transmission dynamics. Temperature, in particular, affects key ecological and biological traits of mosquito vectors such as *Aedes* and *Culex* species, influencing their development, survival, biting behaviour, and the rate of viral replication within the mosquito [9, 10]. One critical epidemiological parameter modulated by temperature is the extrinsic incubation period (EIP), which is the interval required for a virus to disseminate from the mosquito’s midgut, following an infectious blood meal, to the salivary glands, enabling subsequent transmission [11]. Elevated temperatures are generally associated with a shortened EIP, thereby increasing transmission efficiency by reducing the time needed for the mosquito to become infectious [12, 13]. However, excessively high temperatures may increase mosquito mortality, potentially offsetting these gains in transmission potential [14].

The effects of temperature on EIP are not uniform across all systems. Prior research has demonstrated substantial heterogeneity in both the direction and magnitude of these effects, largely depending on the specific vector–virus pairing [15, 16]. However, much of this work has been conducted under constant temperature conditions, which do not accurately reflect natural environments [16]. In reality, mosquitoes are exposed to fluctuating daily temperatures, and there is growing evidence that such diurnal temperature ranges (DTRs) can modulate vector competence. The magnitude of the DTR appears to influence the strength of the response, although findings remain inconsistent [17–19].

The recent unexpected resurgences and re-emergence of many arboviral diseases, including Oropouche virus and Mayaro virus, highlight the urgent need for a better understanding of the transmission dynamics of under-researched viruses [20, 21]. Elucidating the interplay between environmental factors and EIP in both well-known and neglected viruses is essential for improving predictions of arboviral disease dynamics and informing timely public health responses [22, 23].

This systematic review is the first review to comprehensively synthesize existing evidence on the temperature sensitivity of the extrinsic incubation period (EIP) across a wide range of arboviral diseases and mosquito species. By examining how EIP responds to different thermal conditions in diverse vector–virus pairings, the review seeks to clarify the extent to which temperature influences transmission efficiency and the timing of vector infectivity. In doing so, it also aims to identify key patterns, inconsistencies, and research gaps, particularly with regard to understudied pathogens and fluctuating temperature regimes. Ultimately, this work intends to support the development of more accurate models of arboviral transmission dynamics and to inform targeted interventions that can help mitigate the growing global threat posed by these emerging and re-emerging pathogens.

## Materials and methods

### Search

We searched 3 databases (Scopus, Web of Science, and PubMed) on May 3rd, 2025 to retrieve relevant literature. Articles published in any year in English were considered. The full search strategy was as follows: (Arbovir* OR “vector-borne disease” OR “mosquito-borne diseases” OR “aegypti” OR “aedes”) AND (“extrinsic incubation period” OR EIP OR “incubation period” or “viral replication” OR incubation OR “vector competence”). Further references were found through citation searching of included articles.

### Inclusion and Exclusion Criteria

Given the aims of this study, included studies had to 1) consider an arboviral disease affecting human populations, 2) include a mosquito of any species as the vector, 3) study two or more temperatures, 4) contribute novel empirical or narrative evidence of temperature on EIP, transmission, or viral replication, 4) measure the outcome through either detection in the salivary glands or saliva, or through transmission to non-human primates 5) be a published and peer-reviewed article in the English language. We included studies published between 1932 and May 3rd, 2025.

Additionally, we excluded all studies that infected mosquitoes through intrathoracic injection, as injection bypasses the midgut barrier and is thus not an accurate measure of EIP. Reviews, commentaries, duplicates, and non-peer reviewed studies were excluded. Finally, any studies considering infection with possible confounding effects (e.g., Wolbachia infection) were excluded. To ensure consistency between reviewers during title/abstract screening and full-text review, reasons for exclusion were prioritised. This prioritisation can be found in the supplementary index table S1.

### Data Extraction

Data and study characteristics from eligible studies were extracted into an extraction form on Covidence (Covidence systematic review software, Veritas Health Innovation, Melbourne, Australia) accessible to all reviewers. Two-third of all extractions of quantitative data was completed by one reviewer and verified by a second reviewer. Any conflicts were discussed and resolved until consensus was reached. The remaining one-third of extractions were completed by a single reviewer. We extracted general data, including the year of publication, mosquito species, mosquito origin, arboviral disease studied, and viral lineage to give experimental context.

Given the focus of this review to characterise the extrinsic incubation periods, we further extracted data on the transmission efficiency from all studies. Here, transmission efficiency (TE) refers to the proportion of mosquitoes with virus-positive saliva or salivary glands among the total of blood-fed mosquitoes. This data was extracted alongside the day of testing, the viral load given to all mosquitoes, and the mean temperature, and diurnal temperature range of held mosquitoes.

### Statistical and Narrative Analysis

Studies from which the TE could not be extracted, either due to a) non-reporting of the proportions of positive mosquitoes and/or the number of blood-fed mosquitoes b) the reporting of outcomes in figures, from which accurate numbers could not be determined, either from the caption or text, were included in a narrative synthesis of findings.

In order to examine the available data, viral load was standardized across studies. We assumed that tissue culture infectious dose 50 (TCID50) was equivalent to cell culture infectious dose 50 (CCID50), and that plaque-forming units (PFUs) were equivalent to focus-forming units (FFUs). We then assumed a standard conversion of 1 PFU to 0.7 TCID50. Ranges of viral loads, days, or temperatures were set to the mean. Variances of less than 0.5 were considered constant.

Studies with available data were analyzed using linear regression models stratified by vector class and arbovirus grouping, adjusted for viral load and diurnal temperature range, and species-specific variation accounted for. We excluded data in which mosquito cohorts did not become infectious across two or more temperatures, and Dengue data in which the viral load measurement was mouse LD50, given this is not a comparable measure across studies. The linear model otherwise accounted for standardized viral load, where we considered the EIP (time to detection of the pathogen within the mosquito saliva) of each experimental group to be the time of first detection, e.g., the time to which the first mosquito within the experimental group has become infectious.

## Results

Our search obtained 2,250, 805, 262, and 6 articles from the PubMed, Scopus, Web of Science, and citation searching, respectively, for a total of 2,277 studies imported into screening with duplicates removed. Of these, 435 were assessed for inclusion in a full-text review, and 60 studies were included in our final analysis. The full flowchart and reasons for exclusion can be found in fig 1).

**Fig 1.**
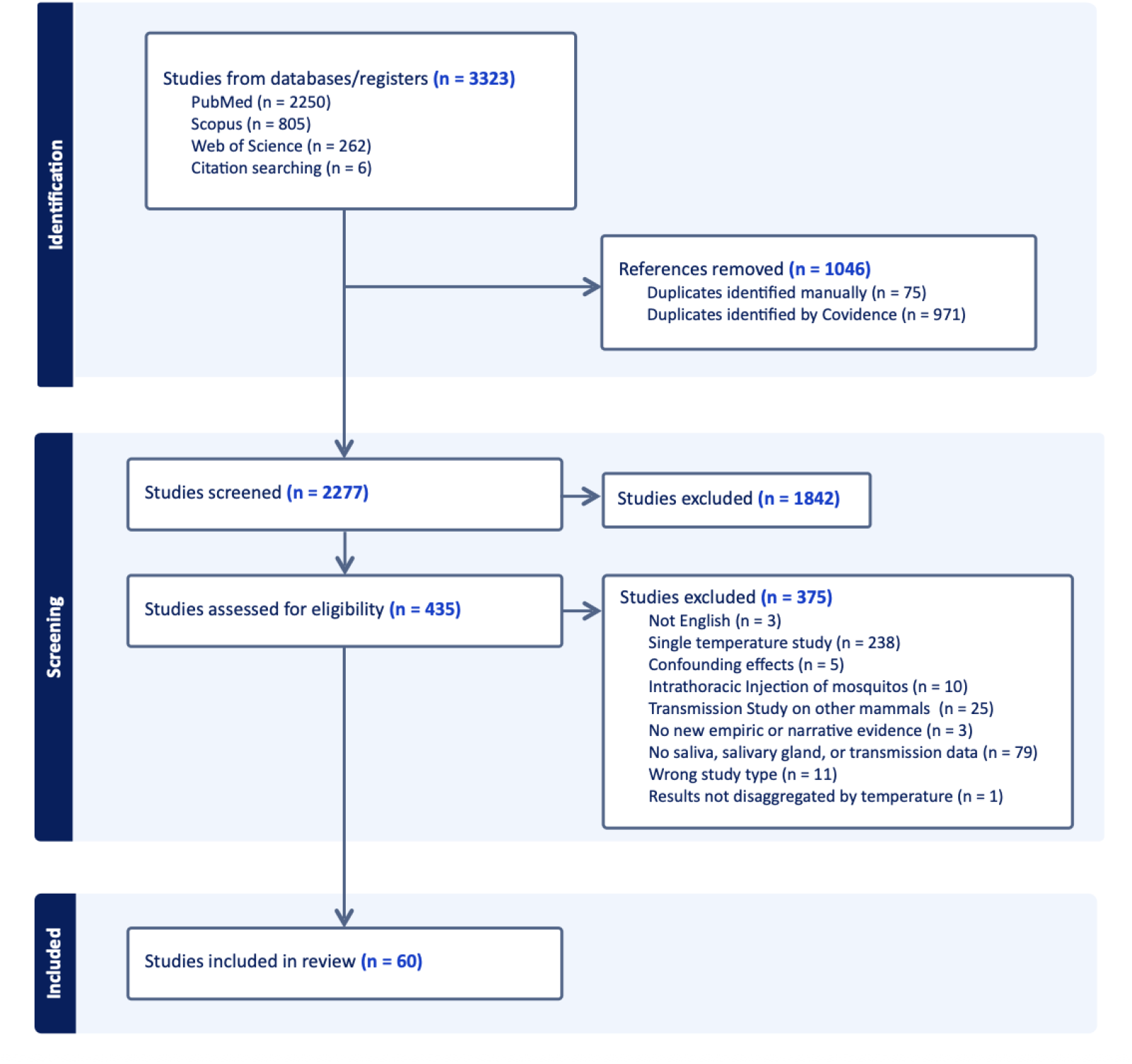
PRISMA Flow Diagram for the Article Selection Process.

### Study Characteristics

Of the 60 included studies, 17 arboviral diseases were studied. Despite a pioneering study in 1932 [24], most research in this field began in the 1980s, and accelerated post-2015, following outbreaks of Zika virus in 2015-2016 (ZIKV) [25] and West Nile Virus (WNV) in 2018 [26]. Many research areas, notably yellow fever virus (YFV), have not show persistent research effort (fig S1).

Of the 60 included studies, data extraction was possible for 48. The 12 narrative studies are further described in the narrative section below. Across the 48 studies for where data extraction was possible, 16 arboviral diseases were studied. The majority of these studied the relationship between temperature and TE for: Zika Virus (ZIKV) [11, 22.9%], West Nile Virus (WNV) [9, 18.8%], and Chikungunya Virus (CHIKV) [7, 14.6%].

This dataset also includes studies across 20 different mosquito species, with the most studied being *Ae. aegypti* (19, 39.6%) and *Ae. albopictus* (15, 31.3%). These mosquitoes originated from 18 different countries, with the most studied in Germany (11, 22.9%) and the United States (10, 20.8%). Only 6 of the included mosquitoes originated from low- or middle-income countries, where the burden of arboviral diseases is highest. An overview of the diseases and mosquito classes studied from each country is depicted in figure 2; here, colour depicts the number of unique experiments conducted across all studies for each mosquito species, disease area, and country (fig 2).

**Fig 2.**
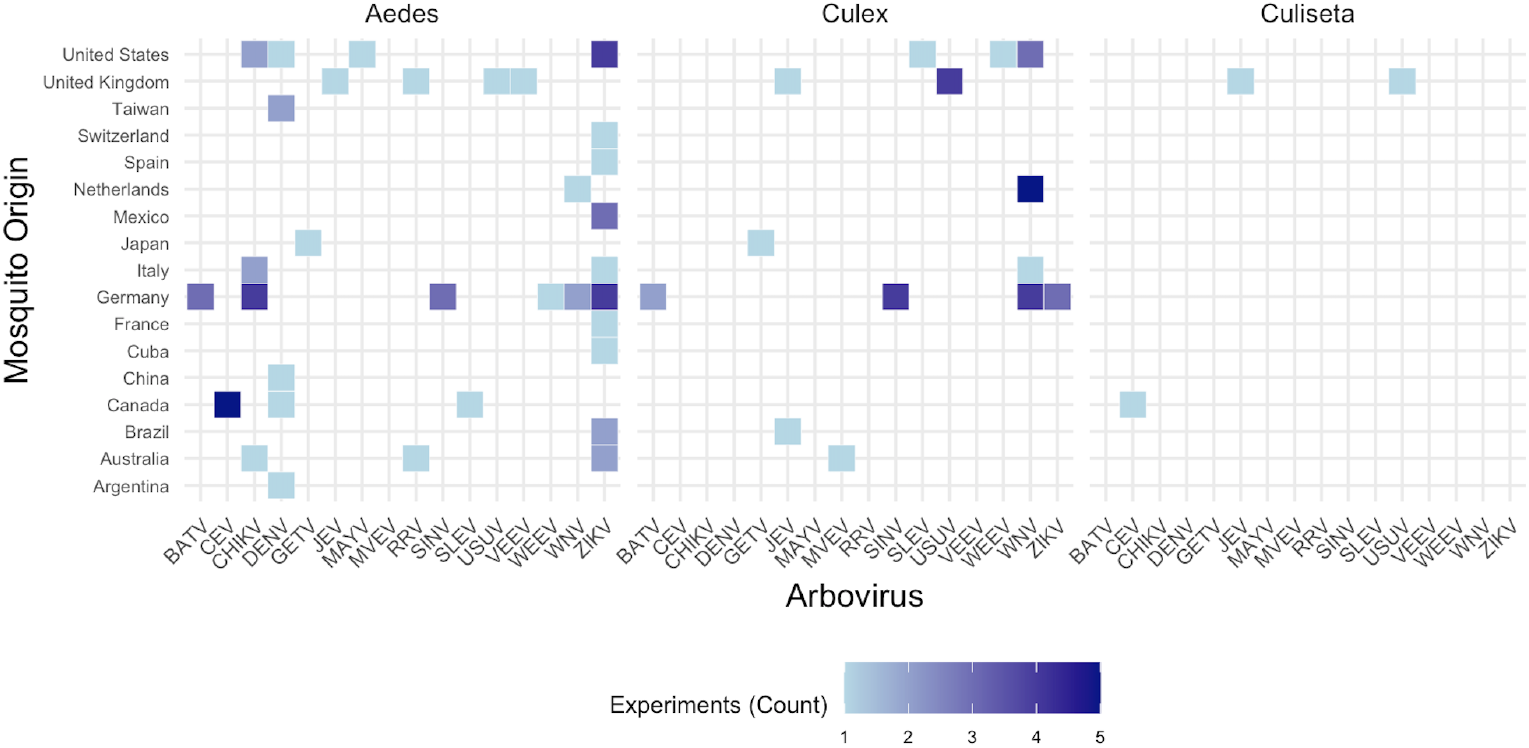
Heatmap displaying the number of experiments conducted for each unique combination of arbovirus, country, and mosquito species (color), with rows representing the countries (y-axis) and columns representing the arboviruses (x-axis). Results are grouped by mosquito class.

37 of the included studies measured transmission efficiency through detection in the saliva; a further 11 in the salivary glands. These studies are described in table 1. Two (4.2%) studies measured transmission through experiments in non-human primates, one on yellow fever and the other on Dengue virus, respectively [24, 27]. These are further described in the section on transmission studies. The dataset containing all final extracted data can be found in SI appendix 1.

**Table 1.**
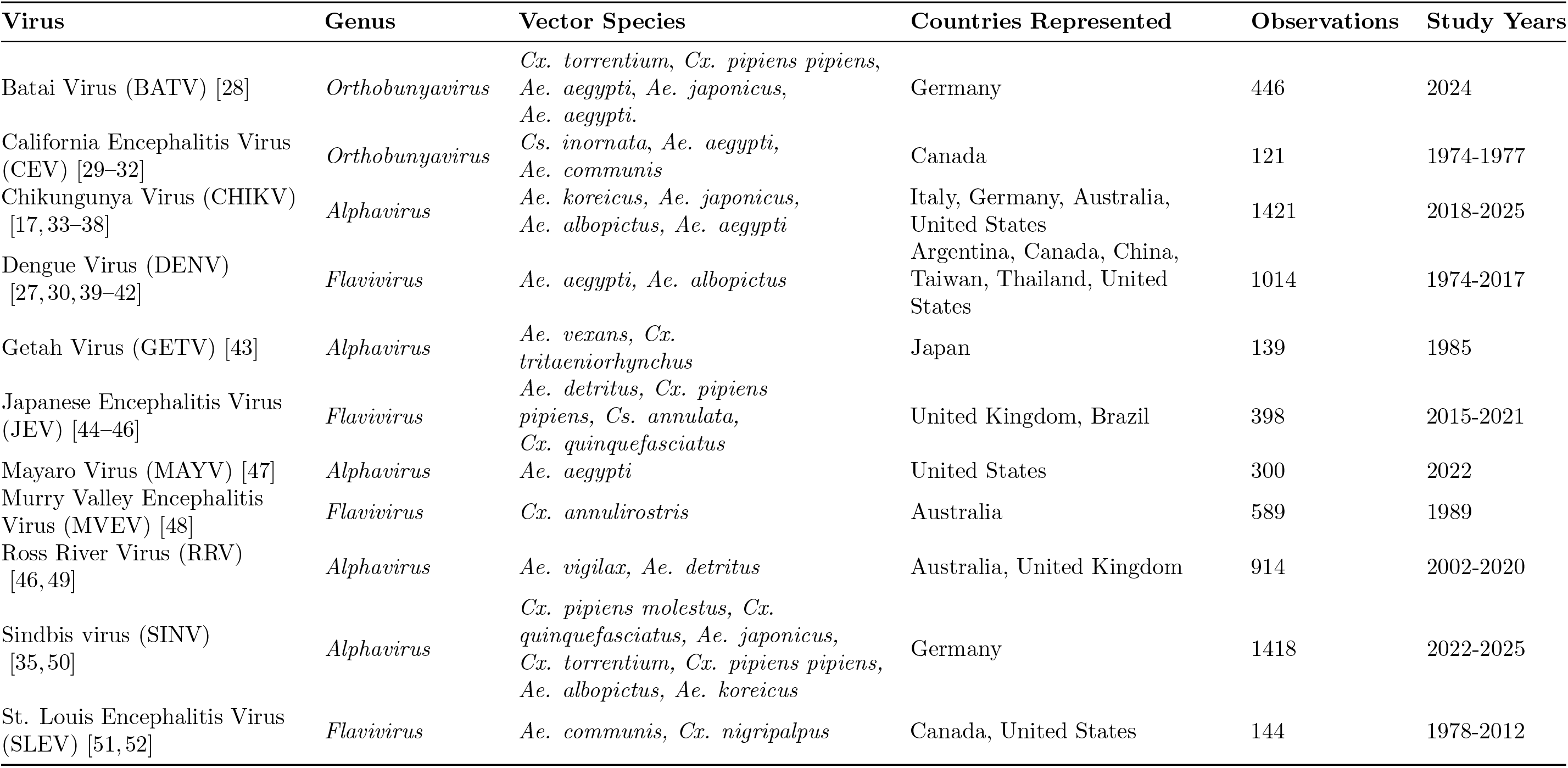
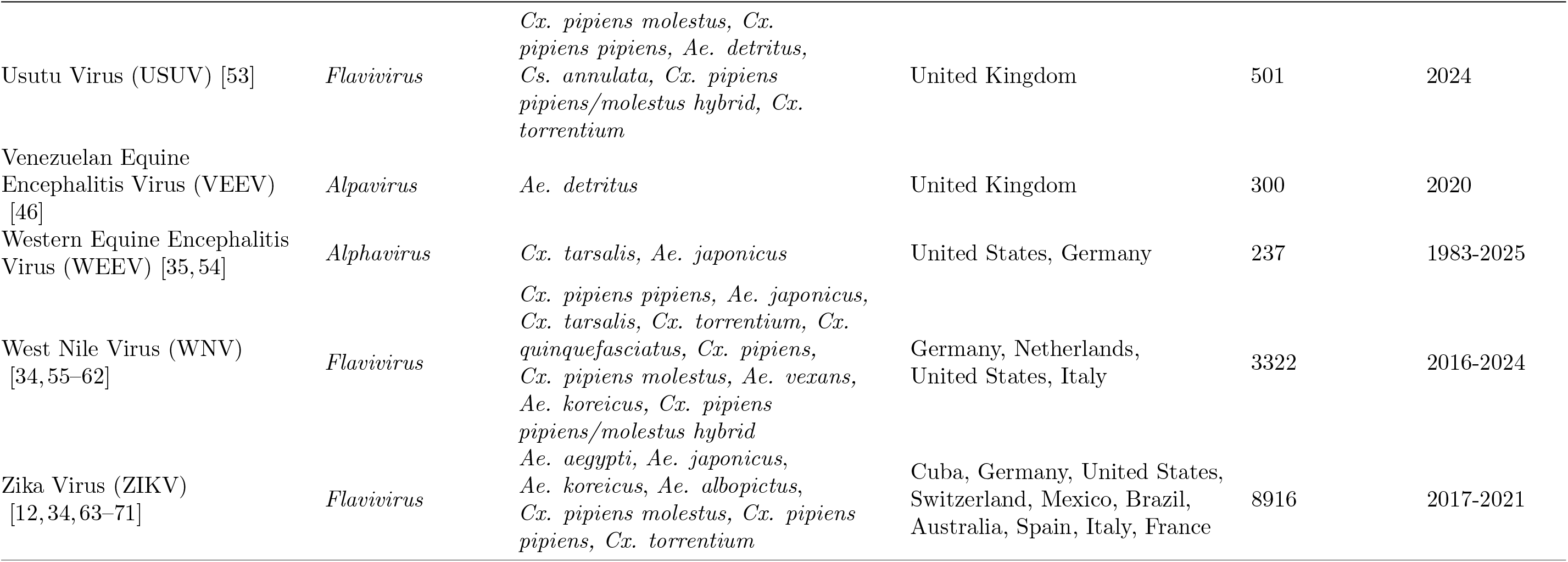
Overview of extracted data for all studies containing transmission efficiency data, as measured by detection in the salivary glands or saliva. Observations refers to the total number of tested mosquitoes, i.e., the denominator of TE.

### Variations in Experimental Design

Experimental approaches across the extracted vector competence studies were highly variable. One major source of this variation was the viral load, i.e., the viral dose that mosquitoes were exposed to. This varied strongly between and within viruses; moreover, the units often varied and required standardization. For example, for Zika virus (ZIKV), experimental doses ranged from 1 × 10^5^ PFU/mL to 4.42 × 10^8^ PFU/mL (in standardized units); despite this, only 1 ZIKV lineage was studied across more than one viral load (American lineage (BRPE243/2015; GenBank KX197192)) [69]. In some vector-virus pairings, such as *Aedes* with St. Louis encephalitis virus (SLEV) or Murray Valley encephalitis virus (MVEV), the observed high transmission efficiencies seen across a single study may in part be due to high viral loads that force infections [48, 51] (figure 3).

**Fig 3.**
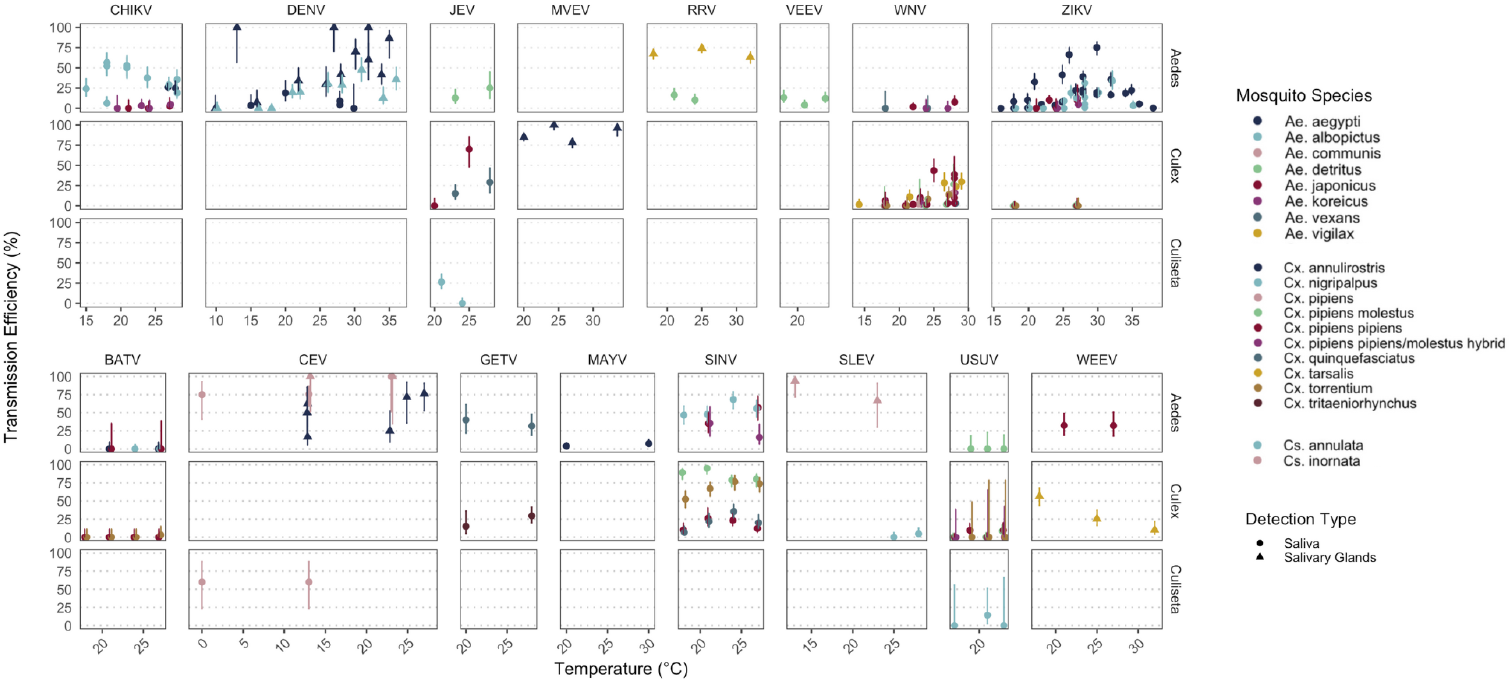
Raw proportion of mosquitoes (grouped by species: color and genus: rows) transmitting (y-axis) the viruses by temperature (x-axis). Each point shows a unique experimental result, i.e. unique viral load, temperature, virus, mosquito species, and study ID; error bars show confidence intervals, calculated using the proportions across one experimental result using the Wilson score interval.

Critically, any studies utilising mouse lethal dose 50 (mouse LD50) have a measure of viral load that cannot be compared to results from other labs or even between experiments of the same lab, as bioassays are highly contextualized; however, in our case, we assume results from the same lab and model animal produced comparable results.

The timing of testing, i.e., the post-exposure window, also varied greatly between studies. For some viruses, like Dengue virus (DENV) and WNV, studies measured broad ranges of data, testing from day 6 to 35 [30], and 5 to 36 days [57], respectively. For viruses with a limited number of studies and/or data, these were often significantly narrowed, i.e., SINV, which only contained measurements on day 5 and 14 [50]. The most commonly observed time points were days 7, 14, and 21.

Finally, the range of temperatures studied across viruses and mosquito species also showed significantly heterogeneity. Fig 3 shows the raw proportion of mosquitoes transmitting virus across temperatures, clustered by mosquito genus and arbovirus, across disease areas. Across the measured temperatures, DENV, Calfornia encephalitis virus (CEV), and ZIKV show the greatest ranges, and Getah virus (GETV), Japanese encepahlitis virus (JEV), and Utusu virus (USUV) show the greatest constraints in data. Across several disease areas, trends can be observed; Chikungunya virus (CHIKV) in *Ae. albopictus*, for example, shows decline in transmission efficiencies at higher temperatures, while DENV shows the reverse. ZIKV, in contrast, appears to have a thermal optimum at moderate temperatures. Variance between vectors can be observed; for example, *Ae. japonicus*, shown to be a poor overall vector of CHIKV, shows no trend. Many arboviruses exhibit no temperature dependency.

### Estimations of EIP

To evaluate the effect of temperature on the EIP, we considered the time to transmission to be the time until the first occurrence of infection in the salivary glands or saliva. We chose this approach following the evidence in Tjaden et al., 2013 [13]: 1) it is most conservative approach, given that it considers the shortest possible EIP and thus does not underestimate risk, and 2) for many vector-virus pairings, the overall transmission efficiency was low, meaning few batches of mosquitoes reached EIP10 or EIP50 thresholds, i.e., the time points in which 10% and 50% of mosquitoes became infectious, respectively, and 3) for many older studies, and by extension, arboviruses, some experimental cohorts consisted of 10 or fewer mosquitoes, which is not otherwise statistically meaningful.

Fig 4 shows the linear regression of time to first occurrence within the vector-virus class, adjusted for mosquito species, viral load and DTR, when varying. Here, a strong heterogeneity of the temperature dependence of the extrinsic incubation period is observed. For several arboviruses, including DENV, CHIKV, WNV, ZIKV, and MVEV, VEEV, the extrinsic incubation period (EIP) declines with increasing temperature, though the strength of this association (i.e., slope) varies across virus–vector combinations. In contrast, no consistent temperature-dependent trend was observed for RRV, CEV, GETV, JEV, SINV, SLEV, USUV, or WEEV, although data for these viruses were often sparse or limited in temperature range (Table 1). Within arboviruses, we did not observe differences in the direction of the temperature effect across mosquito species; however, there were notable differences in intercepts, suggesting species-level variation in baseline EIP. For example, while DENV exhibited similar temperature slopes across species, *Ae. aegypti* consistently showed longer EIPs than *Ae. albopictus* at equivalent temperatures.

**Fig 4.**
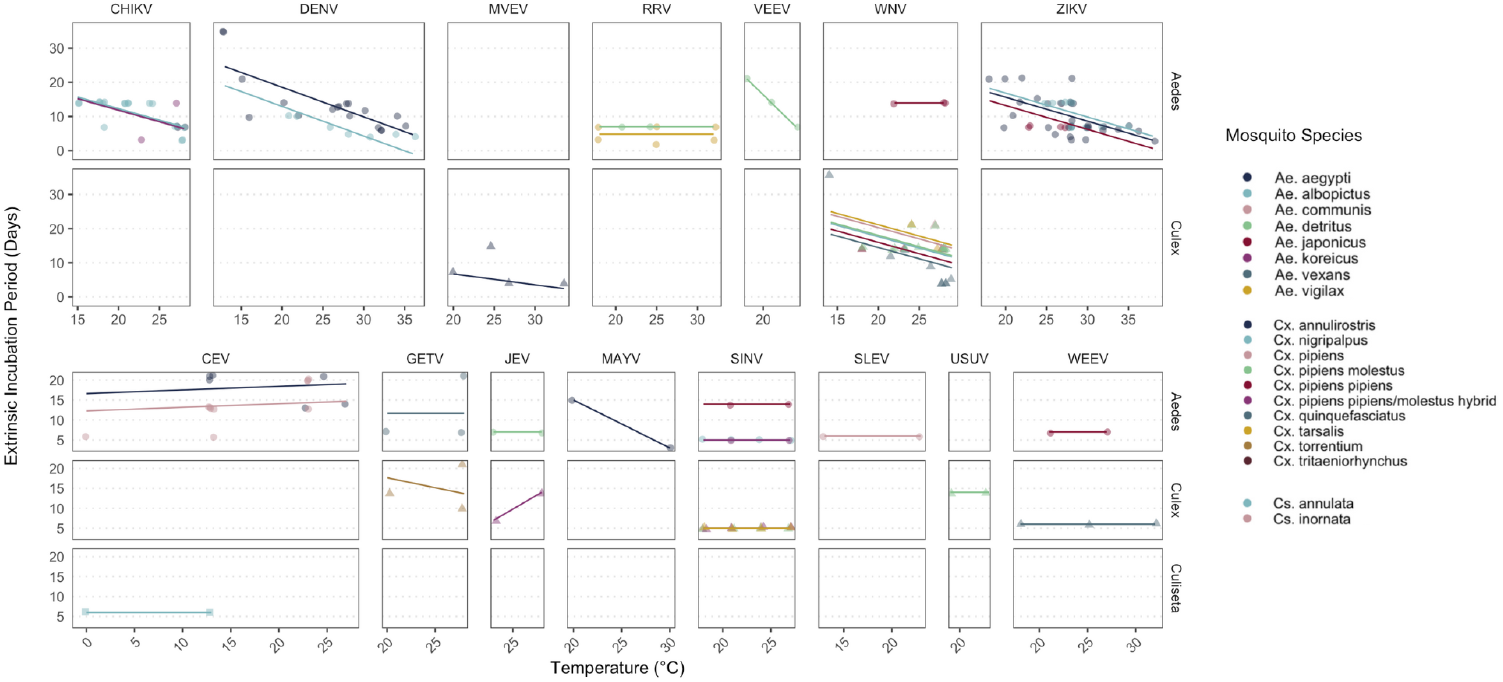
The estimated temperature dependence of the EIP of arboviruses within the dataset. Each point represents the duration until the first observed infection of the salivary gland or saliva, at a given temperature, viral load, DTR, and mosquito species in a given experiment. The solid line displays the linear regression results, with estimates adjusted for viral load and DTR.

### Narrative Results

Across the 12 narratively included studies, temperature was a key determinant of vector competence and transmission efficiency across arbovirus–vector combinations, though its effects varied by species, virus genotype, and experimental conditions.

#### Saliva and Salivary Gland Studies

A total of 10 studies examined the effects of transmission efficiencies and EIP through detection in the salivary glands or saliva.

#### ZIKV

ZIKV transmission increased with temperature across several studies. A study using temperate *Ae. albopictus* and *Ae. detritus* reported a minimum ZIKV transmission threshold between 17–19°C, with higher temperatures generally improving efficiency [72]. In a field-based study from Colombia [73], *Ae. aegypti* showed very low transmission overall, though infection increased at 21 dpi under fluctuating outdoor conditions (24.7–32.9°C). Despite similar average temperatures, daily variation influenced viral titres, indicating that microclimate variability may affect within-host viral dynamics even when mean temperature remains constant.

#### CHIKV and RRV

Elevated temperatures reduced the EIP and increased CHIKV and RRV transmission by *Ae. albopictus*. At 28°C, CHIKV was transmitted in 2 days versus 4 days at 22°C, while RRV was transmitted after 4 days at both temperatures but at higher rates at 28°C [74]. Another study showed the three-way combination of mosquito population, virus strain and temperature further modulated CHIKV transmission [75]. In contrast, no significant temperature effect was seen between 25°C and 30°C at 6 dpi in *Ae. aegypti* from Florida, though sample sizes were low [76]. However, in a field based study from Columbia, researchers found no detectable transmission at either indoor/outdoor fluctuating temperature (mean 28-28.4°C) [73].

#### DENV

Temperature strongly influenced DENV-2 dynamics in *Ae. albopictus*. At 18°C, viral replication remained limited to the midgut. At 23°C and 28°C, dissemination to ovaries and salivary glands occurred by 10 dpi, with significantly higher rates and viral loads at 28°C. At 32°C, the EIP shortened to 5 days with peak transmission. Under fluctuating conditions (28–23–18°C), salivary gland positivity and viral loads were markedly lower than under constant 28°C, suggesting that both mean temperature and stability are critical for efficient transmission [77].

#### USUV

USUV transmission was observed only at 28°C, with an EIP of 7 days and a transmission rate of 38.5%. At 20°C, no transmission was detected. The findings suggest a strong temperature dependency, with warmer conditions overcoming anatomical barriers to transmission [78].

#### WNV

WNV transmission increased sharply with temperature across both Culex and Aedes vectors. In Cx. tarsalis, the both the WN02 (held at 15-32°C) and NY99 (held at 14-30°C) strains saw significant increases in transmission as a function of incubation temperature [79, 80]. In Ae. albopictus, WNV transmission was absent at 20°C but reached 100% at 28°C with a 3-day EIP [78].

#### WEEV and SLEV

One study found the transmission of WEEV and SLEV by Cx. tarsalis increased linearly from 10–30°C, with estimated minimum thresholds of 10.9°C (WEEV) and 14.9°C (SLEV) [81]. Field data showed seasonal transmission peaks during warmer months for SLEV; WEE, in contrast, showed lower susceptibility to transmissions at higher summer temperatures [82].

#### MAYV

Temperature and low relative humidity did not affect Mayaro virus transmission in Ae. aegypti under tested conditions, though exposure to low relative humidity increased mosquito mortality and reduced feeding frequency [83].

#### Transmission Studies

Only two studies examined the relationship between arbovirus transmission in non-human primates and temperature. Both studies demonstrated a clear relationship between temperature and the EIP or transmission efficiency of *Ae. aegypti* for DENV-2 and yellow fever virus (YFV). In Watts et al., DENV-2 was transmitted to monkeys only at *≥* 30^*°*^C, with an EIP of 12 days at 30°C and reduced to 7 days at 32–35°C. No transmission occurred at or below 26°C despite infection rates of 67–95%, indicating a thermal threshold for transmission [27]. Davis et al. similarly showed that YFV EIP shortened with increasing temperature: mosquitoes became infective after 4–6 days at 31–37°C, 8–11 days at 23–25°C, and 18 days at 21°C. No infectivity occurred after 30 days at 18°C unless mosquitoes were later held at warmer temperatures, suggesting partial viral development at low temperatures [24].

## Discussion

Over the last decade, the impact of climate change on health has become one of the most significant global concerns [84]. Research has substantiated that one of these impacts includes the exacerbation of vector-borne disease burden, through both the increased distribution of vectors and population at risk, alongside increased pathogen survival and the shortening of viral replication [85]. In recent years, Dengue, Zika, and Oropouche virus have received significant attention due to major outbreaks or re-emergences [1, 7, 21]; in response, increasing research on the link between temperature and pathogen development within mosquito vectors has been undertaken.

In this study, we sought to characterize existing knowledge on the link between arbovirus pathogen development, the extrinsic incubation period, and climate among mosquito vectors. Our inclusion criteria were broad, spanning both constant and fluctuating temperature regimes, and encompassing lesser-studied viruses and vectors that are often excluded from prior reviews. Despite the increasing evidence that additional climatic factors, such as humidity [83] and rainfall [86], may influence the rate of pathogen development (PDR) and the transmission efficiencies of vector-virus pairings, our review found no studies that focused on links between PDR and climate variables outside of temperature. Thus, this review focuses on synthesizing findings from 60 experimental studies that assess the impact of temperature on arbovirus development within mosquitoes.

Our analysis also brings to light several limitations in the existing evidence base. First, the volume and quality of data are highly uneven across viruses. While DENV, ZIKV, CHIKV, and WNV are well-represented, data for others such as SINV, USUV, GETV, and MVEV remain sparse, with many studies reporting only two temperature points or low overall transmission efficiencies, limiting the ability to infer robust thermal responses. In some cases, data were restricted to salivary gland infection without confirmation of viral presence in saliva, which may underestimate true transmission potential due to salivary gland escape barriers, though prior work suggests a 98.7% concordance between salivary gland infection and vertebrate transmission [87].

Second, methodological inconsistencies remain a barrier to cross-study comparability. There is wide variability in how experiments are conducted, including differences in mosquito rearing conditions, viral doses, and outcome measurement protocols. Critically, there is no standard unit for reporting viral load, despite evidence that dose significantly alters PDR and EIP. Although calls for standardization have been made [88], in practice, these guidelines have not yet been widely implemented across the field. Finally, while we adjusted for viral load and diurnal temperature range (DTR) in our models, DTR showed no major effect, aligning with some previous findings [87]. However, other studies suggest that large temperature fluctuations may affect EIP differently than constant temperatures, suggesting that there may be value in accounting for variation among some vector-virus pairs [19, 89].

Other factors not accounted for may additionally influence the PDR and EIP. Previous research has found that regional differences, influenced by genetic diversity between populations and ecological factors, affect the susceptibility of mosquitoes for arboviral diseases [18, 90–93]. One recent study found that for CHIKV, dissemination efficiency additionally varied based on its K-ancestry [94]. The findings of these studies suggest that vector control strategies should ideally be adapted to the differences based on these regions [95].

Despite these limitations, our synthesis does more than simply confirm existing knowledge. By collating data across all available mosquito–virus combinations, this review provides a clearer and more granular picture of how temperature affects pathogen development. Importantly, we show that even within well-studied arboviruses like DENV, CHIKV, WNV, and ZIKV, the temperature–EIP relationship is not uniform across vector species. While the general pattern of shorter EIPs at higher temperatures holds, the slopes vary, and intercept differences suggest meaningful biological variation, the result of the possible species-specific physiology or genetic factors [91]. These differences are not just academically interesting; they matter for modeling efforts. Most current modeling frameworks assume a single EIP function per virus, often based on data from *Ae. aegypti* alone [14, 96]. Our results demonstrate that this may oversimplify the real-world diversity in transmission potential.

Additionally, the identification of consistent thermal patterns for some viruses and the absence of clear trends for others can help guide future research priorities. Where temperature effects are robust and reproducible (e.g., DENV, ZIKV), these findings can be used to improve parametrization of mechanistic models, estimate uncertainty bounds more realistically, and better inform early warning systems. Conversely, for viruses where data are sparse or trends are inconsistent, our synthesis can help identify where targeted experiments could have the most impact. This is particularly valuable in the context of emerging or neglected arboviruses, where models are urgently needed but empirical data are scarce. Ultimately, our results underscore the need for species- and virus-specific thermal response data, essential for improving the accuracy and applicability of transmission models under future projected climate warming.

## Conclusion

Our results synthesize the effects of temperature across all studied mosquito–arbovirus pairings, offering the most comprehensive overview to date and identifying key patterns alongside critical knowledge gaps. For several major viruses — including CHIKV, ZIKV, DENV, and WNV — we find consistent evidence that higher temperatures can increase transmission efficiency and shorten the extrinsic incubation period, supporting current models of temperature-dependent transmission dynamics. While effects remain unclear for less-studied viruses such as SINV, USUV, and BATV due to limited data, this highlights important opportunities for future research. As climate change continues to alter the distribution and seasonality of vector-borne diseases, our synthesis provides a valuable foundation for improving predictive models. Further experimental work, particularly incorporating under-represented viruses, regions, and additional environmental variables such as humidity, will be essential to refine our understanding and better anticipate future transmission risks.

## Supporting information

**S1 Fig..**
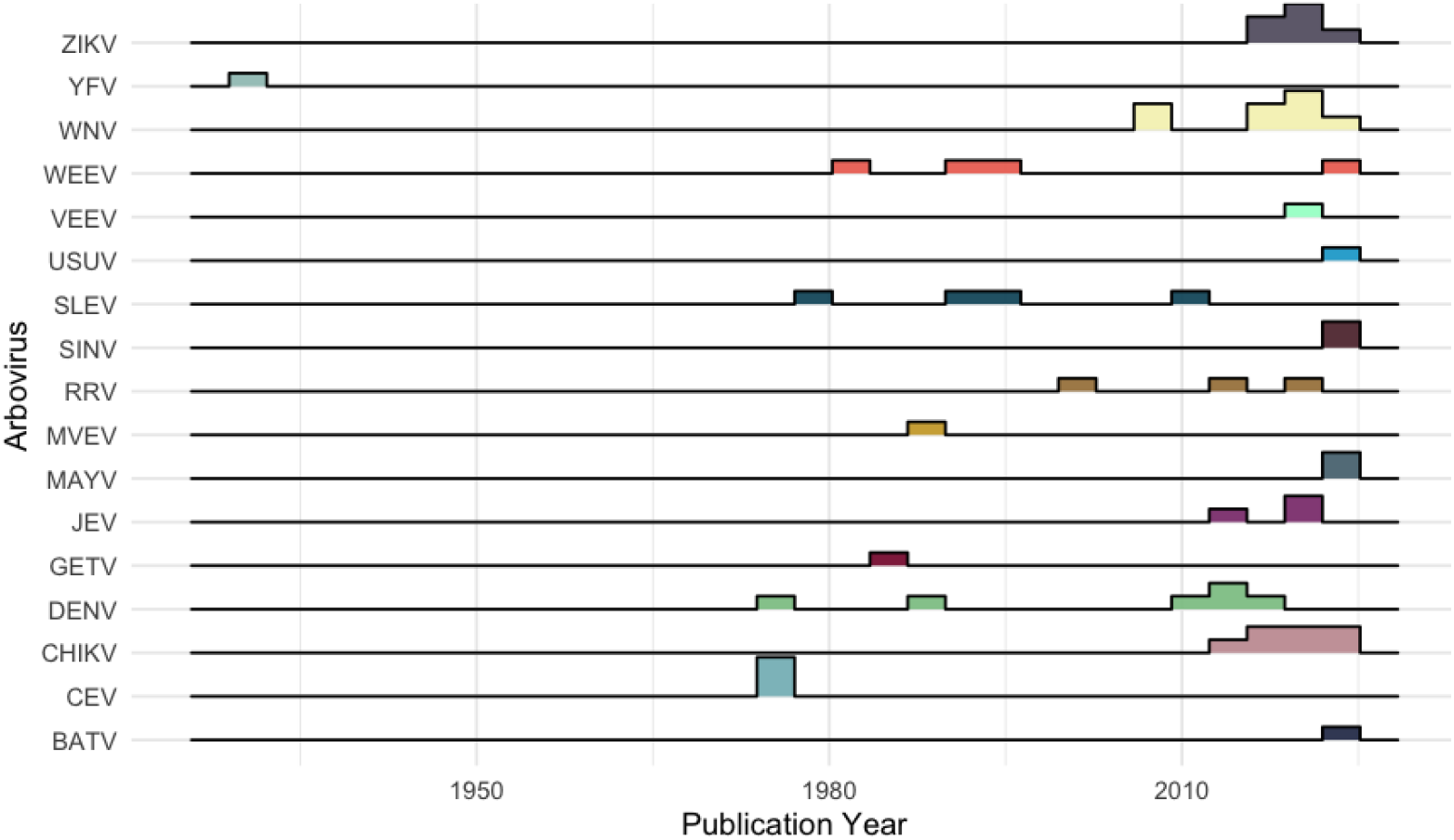
Publication years of all included (narratively and quantitively included) studies. Publications may have included two or more arboviral disease areas and can be represented more than once.

**S1 Table.** Prioritization of study exclusion criteria.

| Item | Reason |
| --- | --- |
| 1 | Not English |
| 2 | Wrong study type |
| 3 | Single temperature study |
| 4 | Intrathoracic injection of mosquitoes |
| 5 | Transmission study on other mammals |
| 6 | No saliva, salivary gland, or transmission data |
| 7 | No new empiric or narrative evidence |
| 8 | Confounding effects |
| 9 | Results not disaggregated by temperature |

**S1 Appendix. Final included dataset.**

## Acknowledgements

AMH and CI acknowledge funding from The Federal Ministry of Research, Technology and Space (BMFTR) through the CLIMADEMIC project (funding code 01LN2210A) within the framework of the Strategy Research for Sustainability (FONA). NK and JP acknowledge funding from the Germany Federal Ministry of Health (BMG) under grant No. 2523DAT400 (project “AI-assisted analysis and visualisation of pandemic situations” — AI-DAVis-PANDEMICS). TH, JBE, LREG, and LW declare no funding. AMH received funding from Gavi, Bill & Melinda Gates Foundation and the Wellcome Trust via the Vaccine Impact Modelling Consortium (VIMC) during the course of the study (grant number INV-034281). All other authors declare no competing interests.

All authors declare no other conflicts of interest.

